# Crop Information Engine and Research Assistant (CIERA) for managing genealogy, phenotypic and genotypic data for breeding programs

**DOI:** 10.1101/439505

**Authors:** Shawn Yates, Martin Lagüe, Ron Knox, Richard Cuthbert, Fran Clarke, John Clarke, Yuefeng Ruan, Jatinder Sangha, Vijai Bhadauria

**Author notes:** Correspondence: Vijai Bhadauria. retired (Fran Clarke).

## Abstract

**Background:** With the advent of next-generation marker platforms and phenomics in crop breeding programs, the volume of both the genotypic and phenotypic data produced has increased exponentially. Often the data remain underutilized if not properly collated, managed and accessed. Effective management of the data is paramount to making sound and timely decision on cross planning in order to accelerate genetic gain (ΔG) in crops for disease resistance, agronomic and end-use quality traits.

**Results:** To address the challenges in managing and efficient utilization of the sheer volume of data generated in a crop breeding program, we developed an electronic information system called the Crop Information Engine and Research Assistant (CIERA). The CIERA, written in Visual Basic, runs on the Microsoft Windows operating system and requires the .Net Framework 4.7 as well as the MySQL Community Server 5.7. The highly intuitive graphical user interface of CIERA includes user-friendly query tools to facilitate the collation of data across relevant phenotypic environments from its phenotypic data management database and can combine that information with the genealogy and genetic data from its genealogy management and genetic data management databases, respectively.

**Conclusions:** Using CIERA, breeders can build a comprehensive profile of germplasm, within a few minutes, to assist them in planning crosses for enhancing genetic gain by selecting superior lines for crosses.

## Background

The success in plant breeding has extensively relied on the availability and access to phenotypic and genotypic information on parental breeding material. However, crop breeding practices have changed dramatically over the past decade. The evolution of marker technology has rapidly increased breeders’ understanding of the germplasm that they develop, and brought a new interest to historical phenotypic and genealogical information. With this advancement in marker technology, the amount of resulting genotypic data can be staggering and often difficult to manage, especially when dealing with next-generation DNA sequencing (NGS) data [1]. In addition, making sense of this genotypic data requires a strong understanding of the phenotype of the germplasm. Handling this massive amount of data is much easier with the use of an electronic information system [2]; although no single system will fulfill all the needs of its users.

In order to achieve trust and acceptance from breeders, a data management system must adapt to the existing practices of the breeding program, be intuitive, and not add significantly to the workload of technicians and researchers. The International Crop Information System (ICIS), was an open-source project led by researchers at the International Rice Research Institute (IRRI) [3], and was considered a good solution for addressing the information challenges faced by breeding programs. ICIS was employed by the wheat breeding programs at the Swift Current Research and Development Centre (SCRDC) from 2001-2015. During this period, the ICIS Genealogy Management System (GMS) and Data Management System (DMS) data models met most of the needs for handling SCRDC breeders’ pedigree and phenotypic data [4].

Researchers found many applications of the ICIS data management capabilities during this time [5, 6]. While ICIS met many of the basic requirements, some of the components were limited or missing, especially in managing genotypic data.

After the ICIS project was discontinued in 2011, an evaluation of existing and upand-coming breeding information systems was undertaken to search for a suitable replacement. The criteria was that a new system would have to meet the data management requirements of all SCRDC breeding programs, have an intuitive interface so that users could adopt it easily, be able to deploy to a local area network (LAN) for multi-user access, and be inexpensive to manage and operate.

Three alternative information systems were evaluated (Table 1): 1) The Integrated Breeding Platform - a Generation Challenge Programme/Bill and Melinda Gates Foundation initiative which uses the Breeding Management System (BMS) for data management purposes, as well as providing a suite of statistical and modelling tools, 2) Agrobase Generation II - a proprietary system from Agronomix Software Inc. which handles plant breeding data, some statistical analysis, reporting, and a host of other features related to a breeding project, and finally, 3) Katmandoo - a system from New South Wales Government Department of Primary Industries which handles phenotypic and genotypic datasets and has some capabilities to handle genealogy data.

**Table 1.** Comparison of existing plant breeding data management systems

| **Softwares** | **LAN**<br>**Install** | **Genealogy**<br>**Management** | **Phenotypic**<br>**Management** | **Genotypic**<br>**Management** | **Support**<br>**Available** | **In-house**<br>**Development**<br>**Opportunity** | **Cost** |
| --- | --- | --- | --- | --- | --- | --- | --- |
| CIERA | Yes | Yes | Yes | Yes | Yes | Yes | Open source (free) |
| ICIS | Yes | Yes | Yes | No | No | Yes | Open source (free) |
| BMS | Yes | Yes | Yes | Limited | Yes | Yes | Annual subscription for<br>some components |
| Katmandoo | No | Limited | Yes | Limited | Limited | Yes | Open source (free) |
| Agrobase | Yes | Yes | Yes | Limited | Yes | No | Multi-user license |

Realizing that the existing ICIS implementation was still the best option for meeting many of the SCRDC breeding program requirements, we decided to develop an improved version, retaining the most used functionality of the predecessor ICIS, and adding new features to address the limitations. This also ensured that bugs and new features would be addressed immediately, and users would not have to wait for a 3^rd^ party to adopt these tasks. To develop this new system, we considered the experiences and user feedback of the users from over a decade. This helped to ensure that the new system would not only meet immediate needs of SCRDC breeding programs, but also be flexible enough to adapt to new breeding practices and evolving data collection technology.

Here we report the development, qualities and use of a new stand-alone electronic information system named the Crop Information Engine and Research Assistant (CIERA), useful for managing breeding data with any crop. This system, in addition to employing ICIS’s approach to handle the genealogy and phenotypic information, has modules to manage genotypic data as well (Fig. 1). CIERA employs a data model that is flexible enough to evolve with the NGS data, besides handling the data generated from Single Sequence Repeat (SSR), Diversity Arrays Technology (DArT), Single-Nucleotide Polymorphism (SNP) and Kompetitive Allele Specific PCR (KASP) marker technology. This data is loaded through the Genetic Dataset Loader and queried through the Data Mining Tool modules.

**Fig. 1.**
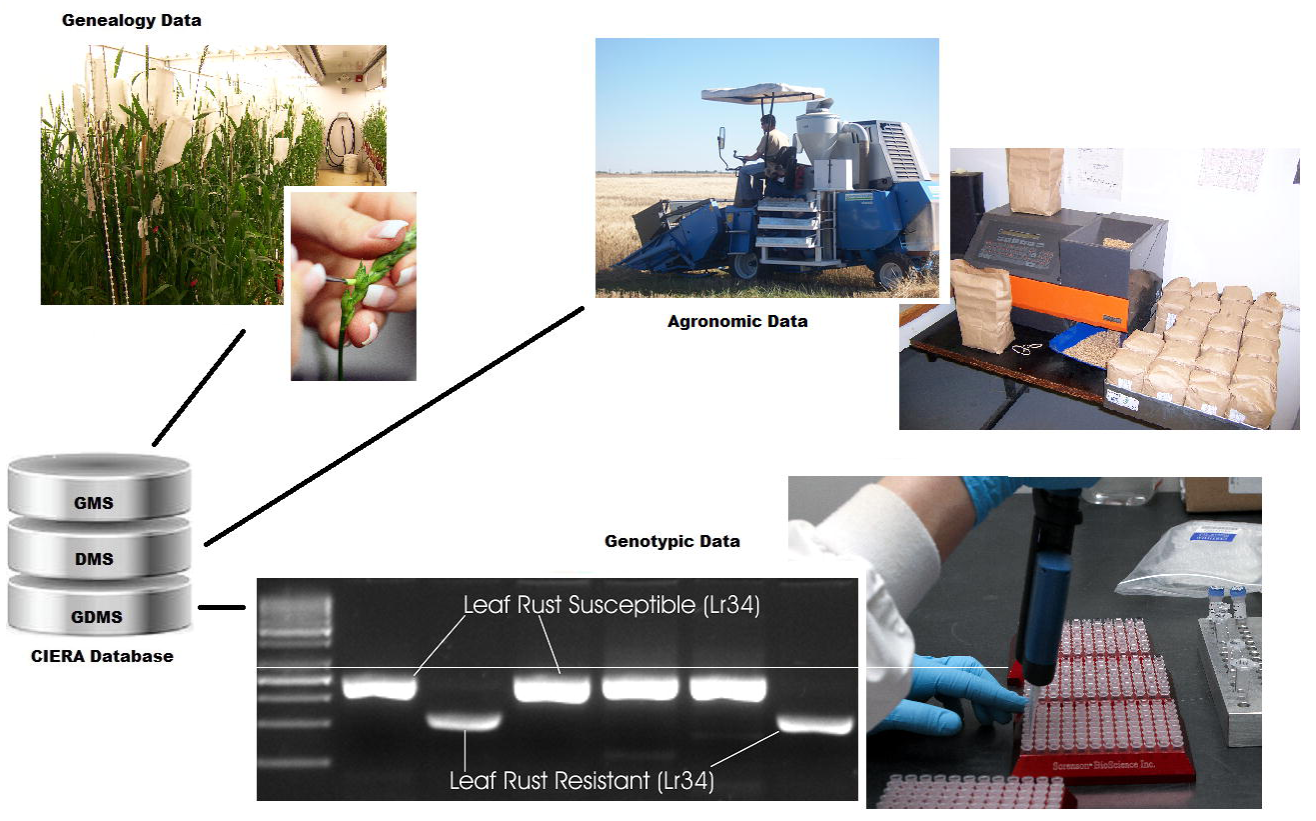
CIERA database system. The data models handle data from phenotypic and genotypic studies, as well as the genealogy of the germplasm

The CIERA standalone version is installed on a client’s machine, but can allow multiple users access to the database on a LAN at the same time. A CIERA web version is under development to port the system to a web interface and move the tools to a web server to improve speed and performance. CIERA, due to ease of use, advanced features of data handling and low cost maintenance, will be useful to the scientific community interested in bringing together all the current and historical phenotypic or genotypic information related to a crop for use in breeding programs.

## Implementation

### System Requirements

The CIERA standalone version currently runs on Microsoft Windows operating system, and requires the .Net Framework 4.7. Due to the potential size of datasets and output, it is recommended that, at minimum, Microsoft Excel 2010 be installed. MySQL Community Server 5.7 is currently the database backend for CIERA. An installer is available for the CIERA launcher and tools on SourceForge (https://sourceforge.net/projects/ciera).

To create a modern and visually enriched interface, Microsoft Windows Presentation Foundation (WPF) (Microsoft 2017) was used for the graphical user interface (GUI) windows, while the rest was developed using Visual Basic (Microsoft 2017).

CIERA has been in use with the breeding programs at SCRDC since 2016, and most issues relate only to hardware limitations particularly on machines with limited RAM when datasets reach millions of rows. No issues with CIERA are reported on high-end machines having at least 16 GB of RAM.

### Databases

In 2012, the International Maize and Wheat Improvement Center (CIMMYT) released a public wheat database to all ICIS/BMS users in the ICIS GMS schema format, containing pedigree information on over six million germplasm from international wheat breeding programs. The information in this database provides wheat breeders a valuable resource about the ancestry of many germplasm lines in the background of the wheat breeding programs, so only minor tweaks were made to the ICIS GMS schema (Fig. 2) to avoid the daunting task of porting the records from this database to a completely new schema. Since users of the legacy ICIS genealogy tools found them quite intuitive, the decision was made to convert the 32-bit tools to a 64-bit platform, but keep the look and feel of their user interface.

**Fig. 2.**
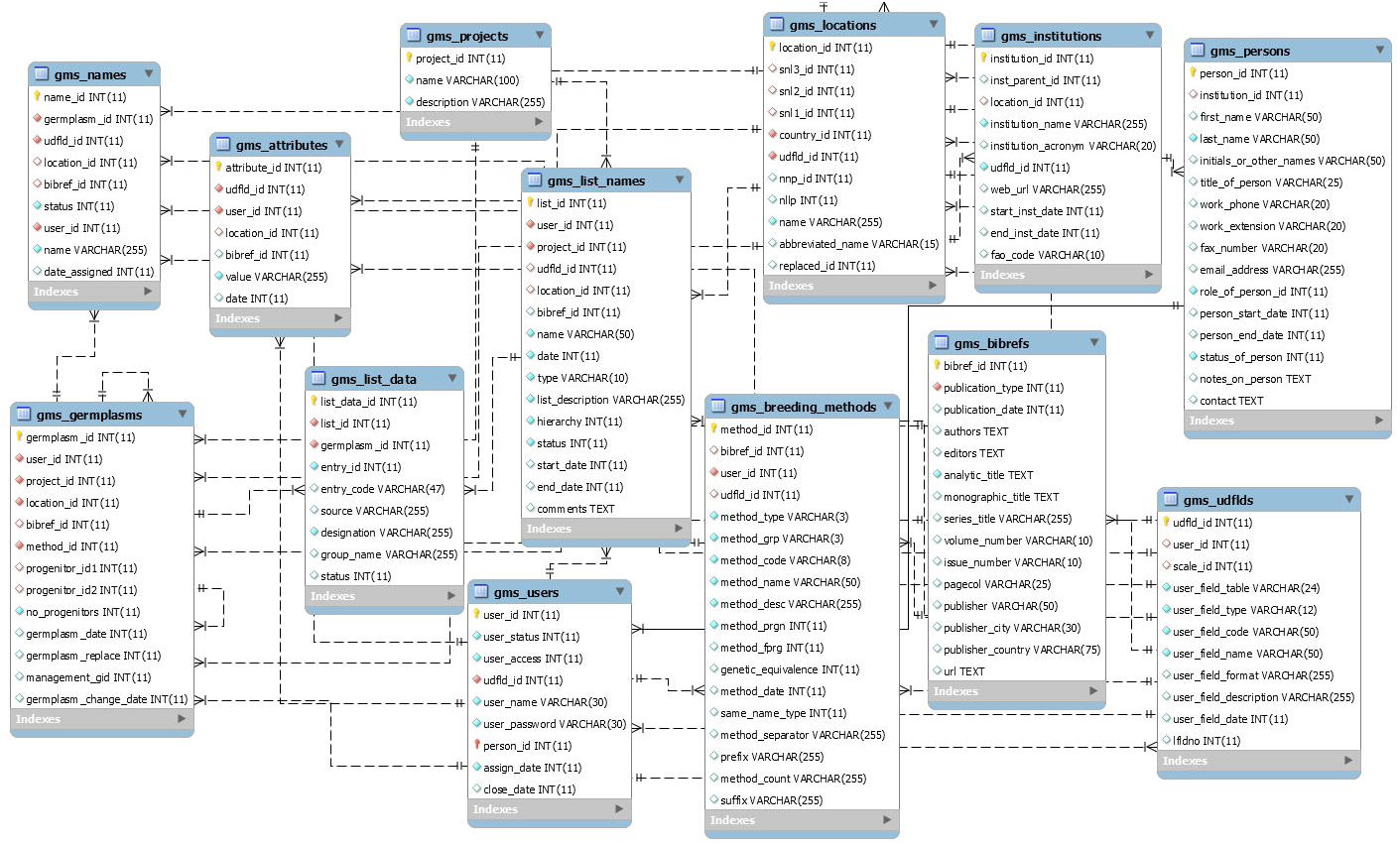
CIERA Genealogy Management System (GMS). All genealogy information on cultivars, passport information and germplasm lists are stored in the GMS

The CIERA DMS is the database schema for storing raw and analyzed phenotypic data at SCRDC, and is based on the ICIS DMS example (Fig. 3). It has proven to accommodate any experimental design, variable or data type required for the breeding programs, and allows breeding programs to define their own variables and nomenclature. This flexibility allows CIERA to adapt to a breeding program, rather than having breeding programs learning new terminology which can lead to mistakes and be cumbersome for staff.

**Fig. 3.**
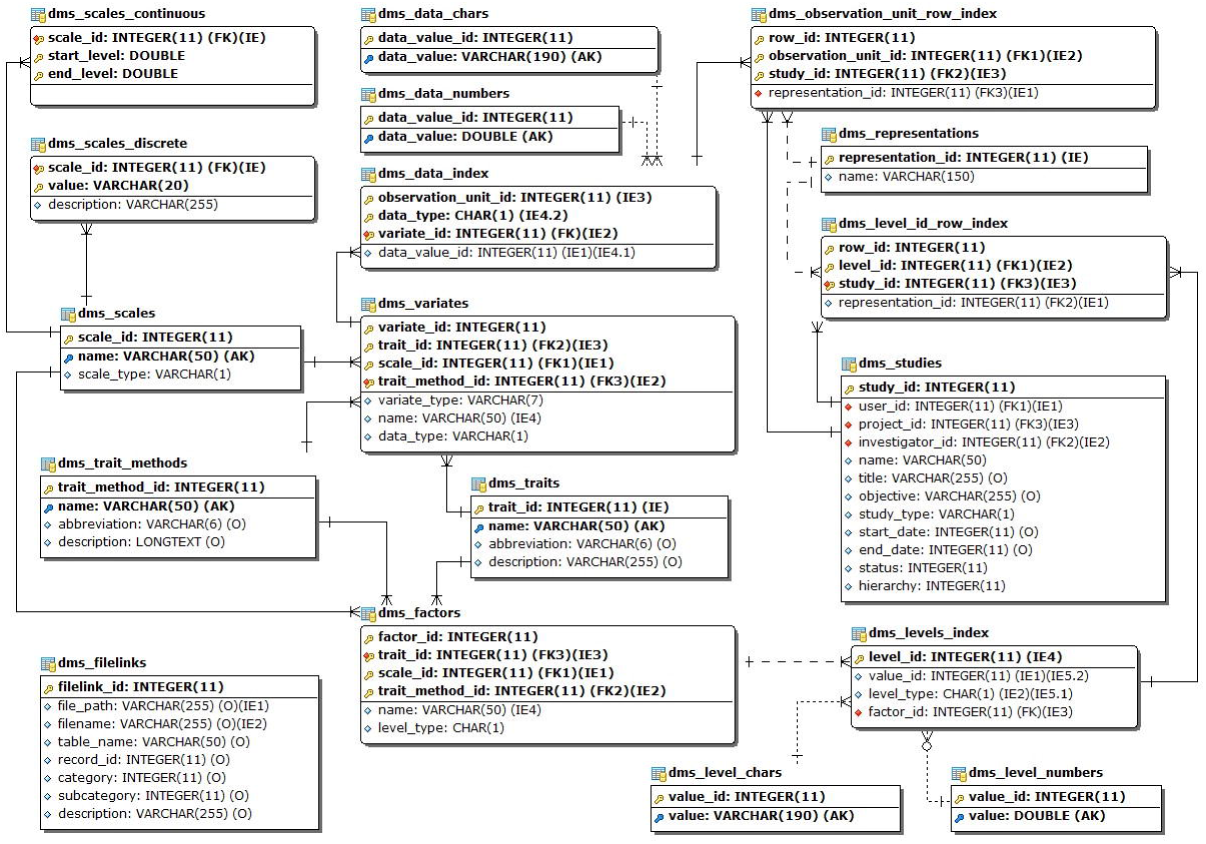
CIERA Data Management System (DMS). Handles the data and study metadata for all types of phenotypic studies, including raw data and summaries

ICIS never had a fully functioning Genotypic Data Management System (GDMS) database schema or tools in production for handling genotypic data, whereas for CIERA, a database was worked out using a modified schema from the GDMS proposed by the Integrated Breeding Platform (IBP) [7]. New objects were added to this GDMS schema, to address missing requirements from the SSR, DArT, SNP and KASP marker datasets that were being used in the breeding programs at the SCRDC (Fig. 4). The schema was designed to handle marker and dataset metadata, alleles (both numeric and character), gene/trait-marker associations and identification of control markers in datasets, quantitative trait locus (QTL) analysis results, and genetic maps. The CIERA GDMS could handle all types of genotypic data by designating the data points as numeric or character, much more useful with NGS technology, generating millions of data points, compared to the few dozen that an SSR dataset might contain. It will be important for researchers to make decisions on which information they want to store in the GDMS, as not all raw reads are useful and these sequences can quickly push the database to an enormous size.

**Fig. 4.**
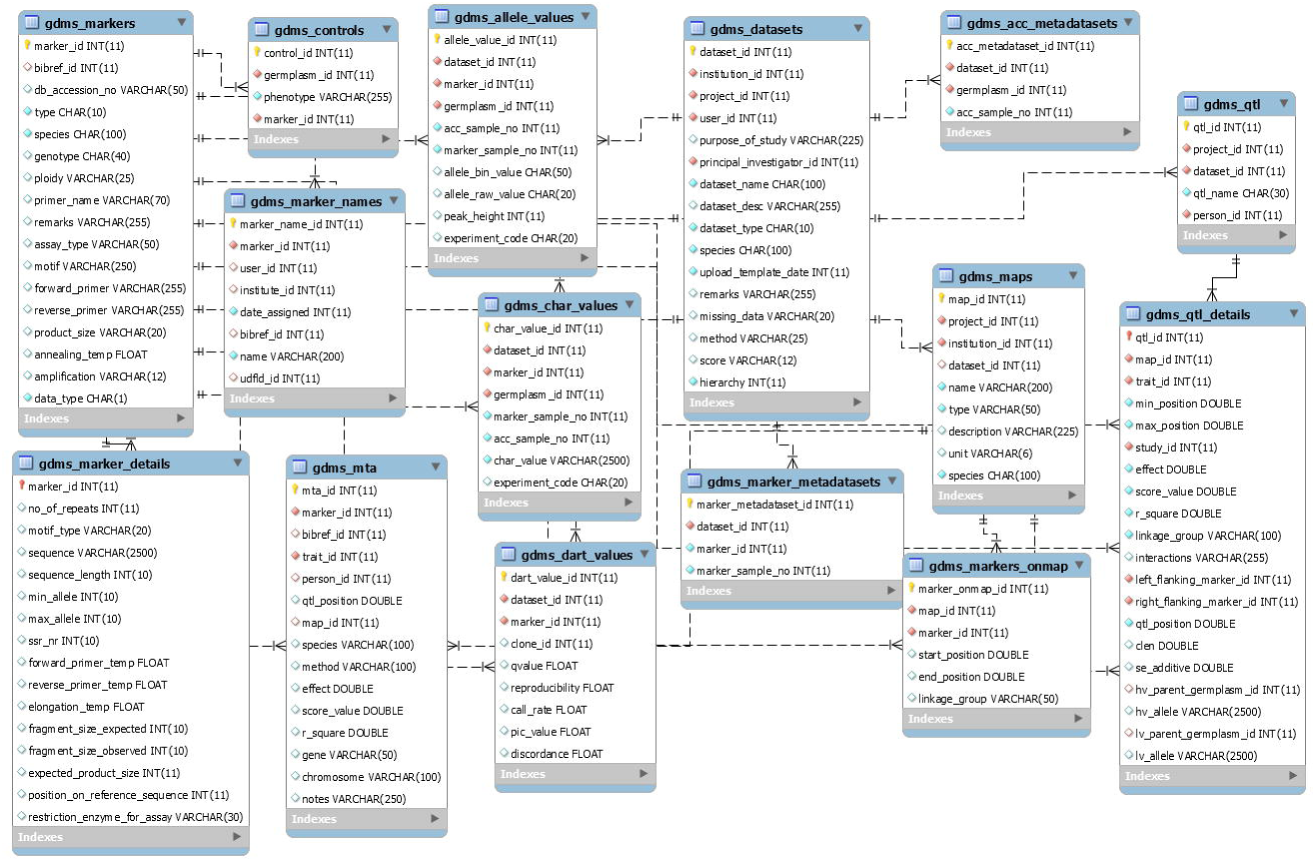
CIERA Genetic Data Management System (GDMS). Schema stores the marker metadata, data from genetic datasets (SNP, DArT, SSR and KASP), QTL information and marker-trait associations

### CIERA Launcher

A CIERA Launcher interface was constructed to facilitate logging into the system and navigating the upload and query tools. The launcher’s interface was developed in consultation with technicians and researchers from various breeding programs to ensure the graphical user interface (GUI) was intuitive.

Users create an account and are assigned an ID within the system that tracks the data they upload into the database and controls the tools and data they have access to, as assigned by the administrator. The menus within the launcher focus on regular tasks that a researcher would query for when looking at breeding program data (Fig. 5). By putting the tools in sub-menus that are related to the same task, the user can easily maneuver between tasks and not be overwhelmed by a long list of available tools.

**Fig. 5.**
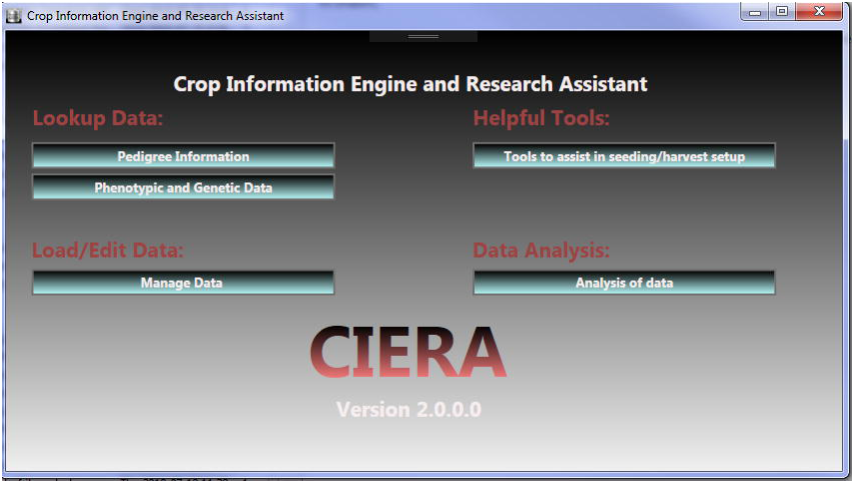
Graphical user interface of the CIERA launcher. The screenshot is of the main menu when starting CIERA

### Genealogy Tools

The tools used for loading and querying pedigree data in the stand-alone version of CIERA are based on the legacy ICIS tools: Genealogy Management System Search (GMS Search), and Set Generation (SetGen). The functionality of these tools is well defined by Portugal et al. [3] and McLaren et al. [4]. By carefully examining the ancestry of potential parents, breeders can make more informed decisions on the sources of disease resistance genes and phenotypic characteristics. This leads to more efficient cross planning and better understanding of the germplasm within the breeding program.

### Genetic Dataset Loader

The Genetic Dataset Loader was developed to facilitate the management of genotypic data. This tool allows users to load, edit and remove genetic data from CIERA.

The first step to loading any genotypic data is ensuring the marker metadata is in the system. The Genetic Dataset Loader handles the loading of marker metadata in two ways: by allowing the user to add information for individual markers, or the user can batch load information on multiple markers at once by filling the marker information into a Microsoft Excel template and loading it. Once in the GDMS, users can edit the marker metadata, link the markers to known phenotypic traits and genes, or remove the marker and all corresponding information (including data from datasets and maps) from the GDMS (Fig. 6).

**Fig. 6.**
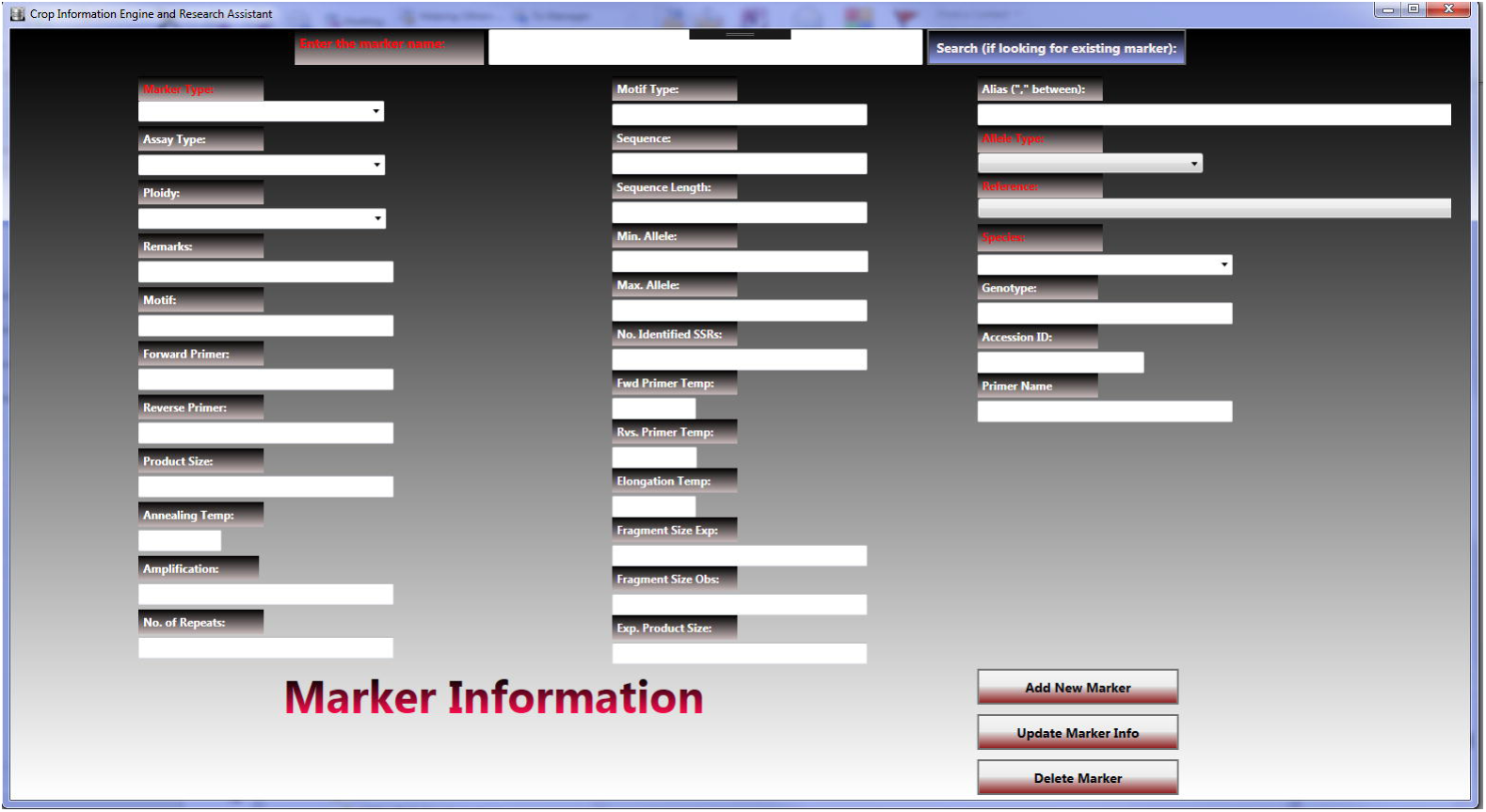
CIERA Genetic Dataset Loader marker information editor: This is a screenshot of the user interface to edit or add marker metadata in CIERA

There are also templates for loading SSR, DArT, KASP and SNP datasets, which can be downloaded from the CIERA SourceForge site. Smaller datasets can be loaded via Microsoft Excel templates, while larger datasets use tab-delimited text files. The Genetic Dataset Loader performs data validation checks prior to importing the dataset into the GDMS, such as making sure the markers and germplasm lines included in the dataset are loaded in CIERA, and that all required fields are filled in.

The Genetic Dataset Loader also manages the importing of genetic maps into the GDMS. The user fills a Microsoft Excel template with map metadata and map information. Upon importing, the tool will again ensure that all markers on the template are already loaded in CIERA, and then move the information to the tables in the database.

Finally, users can add Quantitative Loci Locus, Marker Controls and Marker-Trait Association information through the interface, simply by entering the required metadata on the input screen. This information is used in the genetic data queries to provide more detailed representation of the marker data and assist researchers in making more informed decisions on vast volumes of data.

### Fieldbook Manager

The Fieldbook Manager was developed to facilitate the loading of phenotypic datasets into the DMS. A major problem with managing phenotypic data in any information system is ensuring that the variables are standardized and well-defined within the system. Prior to any loading of phenotypic data, a dictionary of all measured variables from a breeding program should be collected and added to CIERA. This dictionary needs to include the preferred acronym, property (trait) name, scale used, and the method of how the data is obtained, and this should be the starting point for breeding programs wishing to utilize any information system. To assist with this, the Ontology Manager was developed, which allows users to clearly define all aspects of a variable in CIERA.

To import a Microsoft Excel sheet of phenotypic data, the user maps each column heading acronym to a defined variable in the system, or adds a new one. This allows the user to keep acronyms in their spreadsheets that are familiar to them, but ensures at loading time that they are standardized in the DMS. After some validation checks to ensure the data in the worksheet does not have any obvious errors (characters in numeric fields, duplicate rows, etc.), the Fieldbook Manager imports the data to the DMS.

Users can also create their own field books with the Fieldbook Manager, based on a generic template that the SCRDC wheat breeding programs have been using for many years (Fig. 7). The user creates a germplasm list of cultivars, loads a randomization for the test (in a tab-delimited text file, generated from other applications) and some study metadata, along with the variables they wish to include. The tool then generates a standardized Microsoft Excel field book, in plot order, that can be directly sent to collaborators at each breeding test location. The advantage to using these field books is that at loading time, the system will recognize the column headings and directly load the file and all study metadata, saving the time of having to map each column heading.

**Fig. 7.**
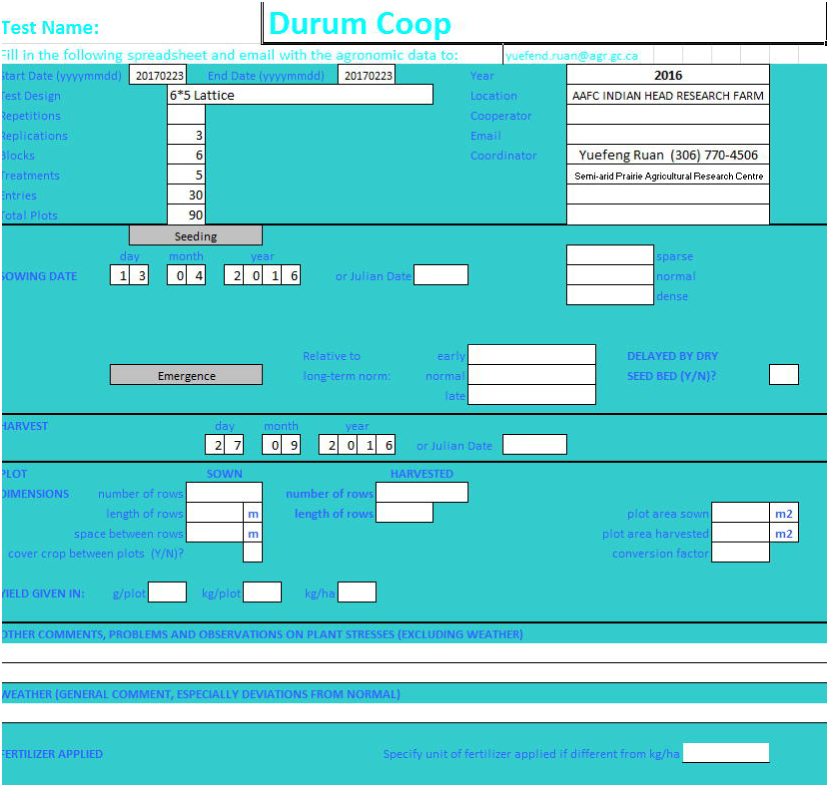
CIERA Field Book cover page: This is an example of the fieldbook cover page that cooperators fill out with study information, such as planting/harvest dates, fertilizer added, and any notes regarding the growing season

The Fieldbook Manager also handles deleting phenotypic studies, should there be any errors to loaded datasets.

### Data Mining Tool

Querying and collation of the genealogy, phenotypic and genotypic data across environments and datasets is a major advantage to using an electronic information system. While using the ICIS implementation, the Data Mining Tool was created at SCRDC using Visual Basic for querying the database. For CIERA, this tool was modified and expanded to incorporate more queries and performance. This tool now has numerous queries (Table 2) to allow researchers to combine phenotypic and/or genotypic data from across datasets and export it to a Microsoft Excel sheet, tab-delimited text file or Extensible Markup Language (XML) file (Fig. 8). The building of a query through a wizard-like approach allows the user to customize the output or filter results not relevant to their needs. This becomes essential when the output from queries is very large. Collation of pertinent phenotypic and genotypic data is paramount when planning crosses and making decisions on which test lines to advance in a breeding program. Making use of the query wizard in CIERA allows breeders to allocate more resources on the decision-making process and less time on the gathering, standardizing and formatting of datasets.

**Table 2.** Data Mining Tool queries, description of query function, output format, and application of the data output

| **Query Name** | **Description** | **Output** | **Use Case** |
| --- | --- | --- | --- |
| Search for data by lines | Enter germplasm identifier individually and retrieve all associated phenotypic and genotypic data; option to add check germplasm lines from the phenotypic datasets | Microsoft Excel<br>Tab-delimited text file<br>XML file | Cross planning; Evaluation of germplasm over environments and time |
| Search for Phenotypic data by study/dataset | Gather whole selected phenotypic datasets | Microsoft Excel<br>Tab-delimited text file<br>XML file | Comparing all germplasm in a particular study, nursery or environment; Evaluating genetic populations; Collating data for reports |
| Search for data on lines in an existing list | Get phenotypic and genotypic data on all germplasm from a list; Get all synonyms for germplasm; Get an overview of all studies the germplasm in the list appear in | Microsoft Excel<br>Tab-delimited text file<br>XML file | Cross planning; Working with international introduction nurseries; Reports of registered variety performance; Quick repeat of queries as new datasets are added |
| Search for genetic dataset or genetic map | Gather whole selected genotypic datasets and/or genetic maps | Microsoft Excel<br>Tab-delimited text file<br>XML file<br>Flapjack .dat and .map files | Evaluating genetic populations; Marker-assisted selection to identify favorable germplasm for crossing |
| Search for information on marker | Search for marker metadata, genotypic data and genetic maps | Microsoft Excel | Follow how germplasm respond to a marker |

**Fig. 8.**
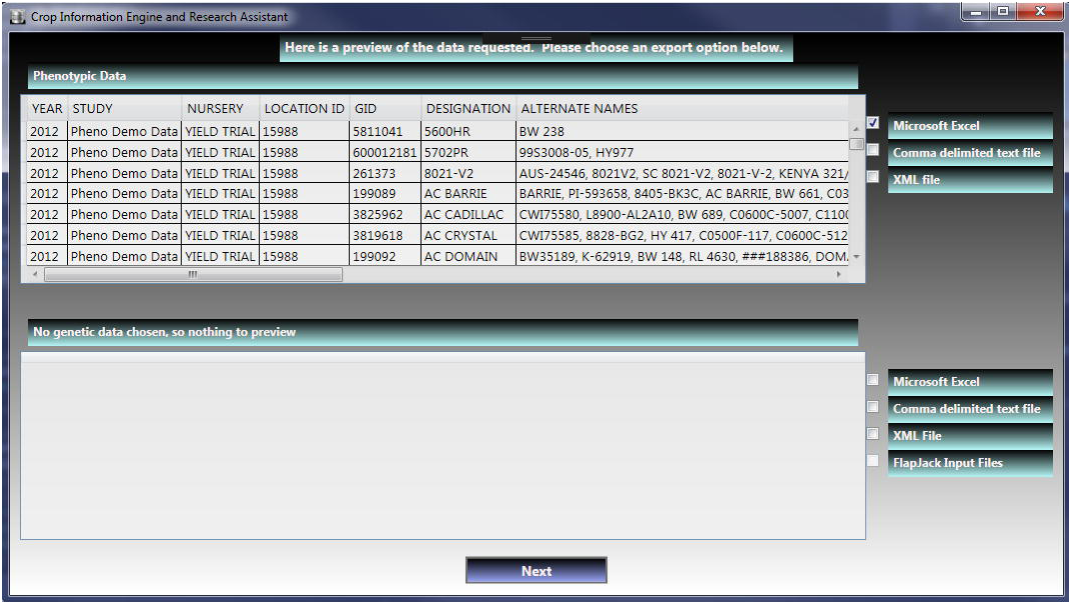
CIERA output preview/export screen: This is a screenshot of the output preview screen, where a user can have a quick look at their query results and choose which format they wish to export to

To assist researchers in evaluating the marker datasets with the visualization tool Flapjack [8], functionality was added to the output routines of the Data Mining Tool to export results from genotypic queries to Flapjack input files.

### Network Deployment

At SCRDC, the CIERA Launcher and tools were installed on users’ machines from a network share via ClickOnce deployment [9], which allows users for easy updating and addresses the problem of having different versions of the software on the computer, by alerting the users to an update upon launching CIERA. A stand-alone installer was also created using the Visual Studio Setup and Deployment project add-in (Microsoft 2017), for users without access to a network, or may require off-line access with the understanding that manual updates will be required in that configuration.

### CIERA Web

In addition to the standalone CIERA, a web-based version is under development. (Fig. S1). This project is using ASPX.net and C# as the programming languages (Microsoft 2017). CIERA web will provide many advantages. The main being the users’ accessibility for CIERA implementation, making the CIERA platform-independent as it will run through a web browser, rather than on the user’s machine. CIERA web also resolves the updating issue, because the tools for CIERA will run off the web server so users will always run the latest version. Another advantage is that, by having the heavy lifting of queries run on the server, users do not need to worry about having high-end workstations for large data sets. Simple devices could be used to access the server (laptops, tablets and even phones), and data could be queried and entered in real time wherever internet is accessible (i.e. greenhouse, laboratory, etc.).

## Results and discussion

The concept for CIERA was to not only store breeding material data, but also to take the next step in assisting researchers to use the data with full potential. The Wheat GMS of CIERA at SCRDC currently stores pedigree and passport information on over six million wheat germplasm. These include crosses, registered varieties, breeder lines, and accessions. Breeders can use this information when planning crosses and can assist in determining the relationships between lines. It also allows breeding programs to track their germplasm samples through CIERA, and link Germplasm IDs with specific bags or envelopes of seed, which is extremely important when tracking seed source of germplasm lines.

The CIERA DMS at SCRDC houses over nine hundred phenotypic studies. These studies include registration yield trials, population evaluations, disease nurseries and quality testing. Where available, both raw data and LS means are stored. The variables are standardized across years and studies, which allows for querying and analyzing the data across environments, and assists breeders in determining candidate germplasm lines to advance through their breeding programs.

As for genotypic data, the CIERA GDMS at SCRDC stores metadata on over forty thousand markers, which include SSR, KASP, SNP and DArT marker types. Close to one hundred genotypic datasets and sixteen genetic maps have also been loaded and are accessible to CIERA wheat users. This data is used for marker assisted selection purpose, but also in marker development and marker validation. As mentioned at the outset, genomics is becoming a major part of modern breeding programs, and NGS technology has been constantly producing massive amounts of genotypic data, which needs proper management and analysis. Deciding which data to store and what format to be used is a challenging task requiring constant evaluation.

CIERA allows the researchers to query the breeding data for genealogy, phenotyping and genotyping information together and simultaneously assemble phenotype and marker profiles of germplasm lines in the breeding program. The Data Mining Tool reduces the time spent on cumbersome efforts of scanning through large spreadsheets filled with thousands of markers on hundreds of germplasm lines to a few minutes, thus saving the resources which would otherwise be required for manual query through datasets and online databases.

### Other Systems

Selecting an information system for breeding program data management is purely a subjective choice; each organization and breeder faces different challenges and employs various approaches to plant breeding. Therefore, a breeder has to assess their program requirements, and choose a system that meets the majority of those needs. Three very capable systems were evaluated as a possible replacement to ICIS at SCRDC in 2015, with the understanding that no one system fits all requirements.

The Breeding Management System (BMS) (https://www.integratedbreeding.net) is the data management component of the IBP. This system not only offers data management tools but also some modelling and statistical tools for breeders, currently not possible with ICIS or CIERA. The BMS is very good at creating phenotypic field books, managing germplasm lists, generating various experimental designs, and loading phenotypic datasets. It also comes with a public wheat database from CIMMYT, which in addition to the genealogy information, has phenotypic and genotypic data as well. However, the feedback from SCRDC users was that the BMS interface was difficult to use, the genealogy tools were not as intuitive as the legacy ICIS tools, and querying phenotypic datasets across studies was difficult, or did not work. Users found they could not search for both phenotypic and genotypic data at the same time, and there were several issues with loading genotypic data. The statistical and modelling components of the BMS were not interesting to SCRDC breeders, as they use other software packages for these purposes. The IBP also requires an annual fee to use some portions of the BMS, with no guarantee that free modules would remain free in the future. Finally, there was a concern that SCRDC issues may not be addressed in a timely manner, as the IBP mandate is to support users in developing countries first.

Agrobase Generation II (https://www.agronomix.com), from Agronomix Software Inc., also offers similar data management features to ICIS and the BMS. It has a suite of tools for performing statistical analysis, tools for printing reports, labels and designing field plans. But Agrobase also requires an annual subscription, and there was concern about how it handles genotypic datasets, specifically NGS data. Being a proprietary system, it would not be possible to develop any new tools to address missing elements or incorporate any in-house tools, as the code is closed to the public. No public wheat database is distributed with Agrobase, so all the pedigree information from the wheat ICIS GMS databases would have to be imported manually through the interface. There was also concern that by using a proprietary system, features may change or be removed without consultation and that issues raised by SCRDC may not be addressed unless they were wide-spread.

Katmandoo (http://katmandoo.org.au/help/Index.htm), from NSW Government Department of Primary Industries, is an open source system that was originally designed to assist breeders in identifying notable germplasm lines through their phenotypic and genotypic data, and subsequently evolved into a breeding data management system. Feedback from SCRDC users regarding Katmandoo was that its interface was not intuitive and required a learning curve to run. There were also concerns that it did not manage genealogy data as well as the ICIS GMS. As with Agrobase, no public wheat database comes with Katmandoo, so the information from the wheat ICIS GMS databases would have to be migrated to the new database or be imported through the interface. Also, it was felt that there were some limitations to being able to load all the different types of phenotypic and genotypic data used in the breeding programs at SCRDC, and that Katmandoo lacked querying capabilities for the data that the Data Mining Tool provided in ICIS.

### Other Crops

In 2016, the Agriculture and Agri-Food Canada (AAFC) Fredericton Research and Development Centre (FRDC) showed interest in CIERA. Their centre focuses on potato breeding, which is a very different type of breeding program from the wheat breeding programs at SCRDC. Since ICIS data models were used for data management in international breeding programs for dozens of different crops in 2011, it was expected that CIERA would also inherit this ability. After some demonstrations and discussions, the breeders at the FRDC did not find any major barriers in having CIERA manage their potato breeding data and this led to collaboration between the Research Centres to share resources. Having a different crop, which utilizes a different type of breeding program, will verify the flexibility of CIERA and corrections will be made should any problem arise, which is one of the major advantages to open source, collaborative development.

### Future developments

The focus going forward will be on completing the CIERA web to replace the standalone version of CIERA. With the large datasets being generated in modern breeding programs, it is necessary to reduce any speed issues while querying or loading data and 64-bit web architecture would provide increased efficiency in computer power [10].

In addition to the CIERA web development, another focus is to integrate many of the independent in-house tools that technicians at SCRDC use for seeding and harvest tasks, similar to the suite of tools in Agrobase. These tools will make use of the information stored in the database instead of exporting and importing input files. This will increase the efficiency of the breeding program activities and also reduce errors. Breeding programs also use many 3^rd^ Party proprietary and open source software tools to perform various tasks, analysis, visualization, reports and printing labels. While it is not prudent to develop new tools to replace existing 3^rd^ Party software that already meets the required needs of users, input files can be prepared if the software can make use of the data stored in CIERA. Manually creating these input files can be time-consuming, and there is always concern for errors during preparation.

## Conclusions

CIERA was built upon the knowledge and experiences gained from collaborating for several years with an excellent development team at IRRI and the feedback from researchers, technicians and computer science staff from various organizations and breeding programs. The goal for CIERA was to create a system that facilitates the researchers working with modern breeding program data, as this data has become too cumbersome to handle solely with spreadsheets. CIERA is highly efficient as it possess all the abilities for data management for standardization, storage, and computing power. The lessons learned during the collaboration between SCRDC and FRDC, each with very different breeding programs, will provide a path to expanding CIERA to other crops and research centres going forward. Future developments will focus on finding ways to improve the usage of the data and make it more informative for researchers.

## Availability and requirements

**Project name:** Crop Information Engine and Research Assistant (CIERA)

**Project home page:** https://sourceforge.net/projects/ciera

**Operating System:** Microsoft Windows

**Programming Language:** Visual Basic

**Other requirements:** MySQL .NET Connector, MySQL Community Server 5.7, Microsoft Excel 2010 (minimum)

**License:** GNU General Public License, Version 3.

**Any restrictions to use by non-academics:** No restrictions

**List of abbreviations**
AAFC: Agriculture and Agri-Food Canada
BMS: Breeding Management System
CIMMYT: International Maize and Wheat Improvement Center
DArT: Diversity Arrays Technology
DMS: Data Management System (phenotypic)
FDRC: Fredericton Research and Development Centre
GDMS: Genotypic Data Management System
GMS: Genealogy Management System
GUI: Graphical User Interface
IBP: Integrated Breeding Platform
ICIS: International Crop Information System
IRRI: International Rice Research Institute
KASP: Kompetitive Allele Specific PCR
LAN: Local Area Network
NGS: Next-generation sequencing
QTL: Quantitative Trait Locus
RAM: Random Access Memory
SCRDC: Swift Current Research and Development Centre
SNP: Single-Nucleotide Polymorphism
SSR: Single Sequence Repeat
WPF: Microsoft Windows Presentation Framework
XML: Extensible Markup Language

## Acknowledgements

The development of CIERA was funded by the Canadian National Wheat Cluster program. We would like to thank the technical and development assistance of Thomas MacGregor, Jonathon Florek, Jordan LeBlanc and Calvin Blacquier. We would also like to acknowledge the excellent work put in by the ICIS development team lead by Dr. Graham McLaren, Dr. Richard Bruskiewich, Arllet Portugal and Dr. Thomas Metz, formerly at the International Rice Research Institute (IRRI).

## Ethics approval and consent to participate

Not applicable.

## Consent for publication

Not applicable.

## Availability of data and materials

- A demonstration MySQL database, filled with data from templates containing various phenotypic and genotypic data is available on the CIERA sourceforge.net repository, https://sourceforge.net/projects/ciera/files/Demonstration%20Files
- A demonstration tutorial is also available on the project wiki on sourceforge.net,. https://sourceforge.net/p/ciera/wiki/Demonstration

## Author contributions

SY, VB and ML developed the CIERA; RK, FC, JC, RC and YR provided input, testing and feedback for CIERA; SY and VB wrote the manuscript; and JS provided suggestions on the manuscript.

## Supplementary information

**Fig. S1.** Web version of CIERA: This is a screenshot of the CIERA web interface, which is still under construction.

